# Discovery of Expression-Governing Residues in Proteins

**DOI:** 10.1101/2025.01.06.631498

**Authors:** Fan Jiang, Mingchen Li, Banghao Wu, Liang Zhang, Bozitao Zhong, Yuanxi Yu, Liang Hong

**Affiliations:** School of Physics and Astronomy, Shanghai Jiao Tong University, Shanghai, 200240, China; Institute of Natural Sciences, Shanghai Jiao Tong University, Shanghai, 200240, China; School of Information Science and Engineering, East China University of Science and Technology, Shanghai, 200237, China; Shanghai Artificial Intelligence Laboratory, Shanghai, 200232, China; School of Life Sciences and Biotechnology, Shanghai Jiao Tong University, Shanghai, 200240, China; Zhangjiang Institute for Advanced Study, Shanghai Jiao Tong University, Shanghai, 201203, China; Shanghai National Centre for Applied Mathematics (SJTU Center), MOE-LSC, Shanghai Jiao Tong University, Shanghai, 200240, China

## Abstract

Understanding how amino acids influence protein expression is crucial for advancements in biotechnology and synthetic biology. In this study, we introduce Venus-TIGER, a deep learning model designed to accurately identify amino acids critical for expression. By constructing a two-dimensional matrix that links model representations to experimental fitness, Venus-TIGER achieves improved predictive accuracy and enhanced extrapolation capability. We validated our approach on both public deep mutational scanning datasets and low-throughput experimental datasets, demonstrating notable performance compared to traditional methods. Venus-TIGER exhibits robust transferability in zero-shot predicting scenarios and enhanced predictive performance in few-shot learning, even with limited experimental data. This capability is particularly valuable for protein design aimed at enhancing expression, where generating large datasets can be costly and time-consuming. Additionally, we conducted a statistical analysis to identify expression-associated features, such as sequence and structural preferences, distinguishing between those linked to high and low expression. Our investigation also revealed a correlation among stability, activity and expression, providing insight into their interconnected roles and underlying mechanisms.

## 1 Introduction

Protein expression play a pivotal role in cellular function [1] and biotechnology applications [2], directly impacting biological processes and serving as key economic indicators in industrial production by determining process efficiency and product yield [3–5]. Furthermore, mutations affecting protein expression can lead to various diseases, highlighting its clinical significance [6, 7]. Therefore, accurate prediction and manipulation of protein expression have become increasingly important for protein engineering and therapeutic development.

However, it remains difficult to directly infer expression from sequence information alone. In protein engineering, while rational design [8, 9] and semi-rational [10] design approaches have shown remarkable success in improving protein stability and activity [11], they have limited capability in modulating expression. Although directed evolution can potentially achieve expression optimization, it requires extensive experimental screening [2]. Computational methods leveraging homologous sequence information can reduce the amount of experimental data required for protein optimization [12–14]. However, these approaches typically depend on multiple sequence alignments—constraints that limit their applicability to proteins lacking extensive homologs—and have not yet demonstrated substantial utility in protein expression prediction tasks.

Recent advances in protein language models (PLMs) offer a promising alternative. Trained on diverse protein evolutionary sequences, PLMs can learn informative biological representations, thereby enabling advances in protein structure prediction and functional annotation [15–18]. Leveraging PLMs to guide directed evolution has rapidly become a mainstream methodology, with demonstrated success in optimizing fitness parameters such as enzymatic activity, protein stability, and antibody affinity [19–21]. Likewise, emerging applications in de novo protein design are harnessing PLMs to generate novel proteins, expanding the scope of computational tools for protein engineering [22–24]. Despite these advancements, the transferability of PLMs remains a limiting factor [15, 18]. Although PLMs demonstrate sophisticated sequence representation capabilities, their application in optimizing protein expression has yet to yield substantial practical success. Addressing this challenge is essential for fully realizing the potential of PLMs in both directed evolution and de novo protein design.

In this study, we present Venus-TIGER (Tool for Identification of Governing Expression Residues), a protein expression prediction model designed to infer the effects of specific mutations on expression. Unlike conventional strategies that combine a PLM with a top-layer regression model, Venus-TIGER learns the protein expression landscape by comparing a PLM-derived pseudo-matrix with an experimentally derived matrix. This approach enables simultaneous modeling of the entire expression landscape while pinpointing mutations that enhance expression, thereby reducing training costs and improving the model’s transferability. Our approach demonstrates robust performance across public deep mutational scanning (DMS) datasets and exhibits strong transferability to low-throughput experimental datasets, as validated in two representative cases: T7 RNA polymerase (T7 RNAP) and the variable domain of the heavy chain of a nano-antibody (VHH). Through comprehensive statistical analysis, we identified common patterns and regulatory mechanisms by which mutations influence protein expression across different proteins. Furthermore, we extended our analysis to investigate the correlations among expression, stability, and activity. This integrated approach not only advances our understanding of protein expression determinants but also provides a practical tool for protein engineering applications.

## 2 Results

### 2.1 Development of a Landscape-Based Deep Learning Framework for Discovering Expression-Governing Residues

#### 2.1.1 Describing Single-site Mutations’ Expression Level in a Matrix

The effects of single-site mutations in protein sequences on expression can be described using a real-value matrix [25]. The columns in the matrix represent sequence positions; the rows are possible amino acid variants at each position, and the matrix elements represent the expression of corresponding mutants. (Fig. 1c). We refer to this matrix as the “expression landscape”. However, achieving a comprehensive expression landscape is challenging because it requires experimentally measuring all possible single-site mutations. Consequently, this matrix typically only contains values for a limited portion in most instances.

**Fig. 1.**
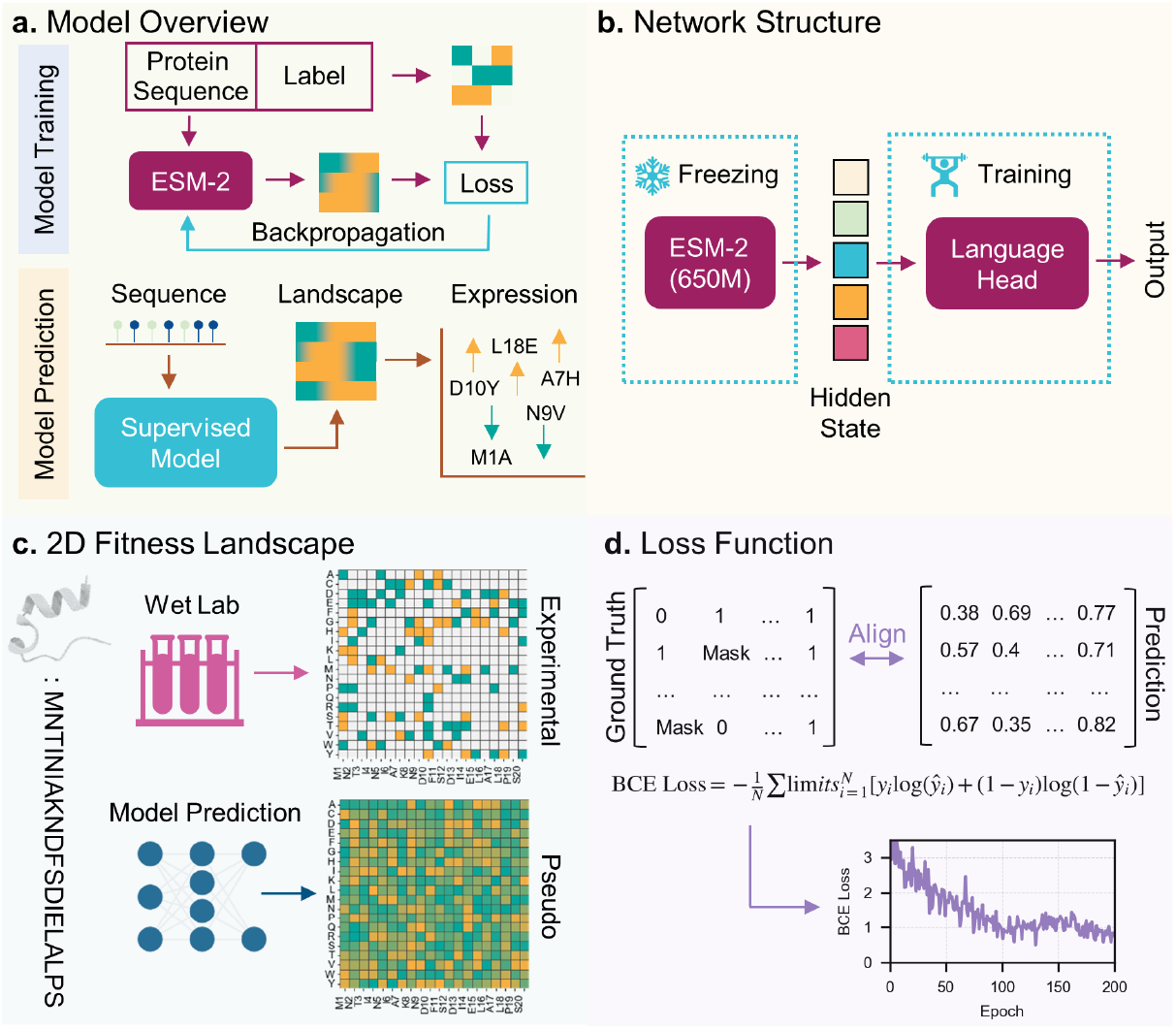
Overview of Venus-TIGER framework. **a.** Model overview: (i) Model training: protein sequences are encoded by pre-trained language models and trained with expression labels; (ii) Model prediction: sequence is processed through the trained model to predict expression for all mutants. **b**. Network structure showing the frozen ESM-2 feature extraction layers and the trainable language head for prediction generation. **c**. Comparison between experimental and predicted 2D fitness landscapes representing mutation effects on protein expression. **d**. Loss function: Implementation of landscape-based BCE loss calculation, applied only to unmasked positions in the training data.

#### 2.1.2 Learning to Align PLM Zero-Shot Prediction with Expression Landscape

PLMs can predict the mutation effect by calculating the odds between the likelihood of the mutated amino acid and that of the wild-type amino acid [16]. However, such predictions are “undirected” since they are solely based on the likelihood learned from the protein sequences in the training set and not targeted towards any specific physical and chemical properties.

To convert this “undirected” prediction into a “directed” one, we propose a method to align the predicted landscape of protein language models with the expression landscape by training on mutation expression level datasets. The overview of our model is shown in Fig.1 and the details of our model are shown in the Method Section 4.1. The architecture comprises two key components: a frozen feature extraction backbone from ESM-2 (650M version^1^) [15] and a trainable language head (Fig. 1b). The backbone and language head are all inherited from the original ESM-2, with no additional parameters added. The framework implements an innovative sequence-to-landscape training paradigm that fundamentally differs from conventional sequence-to-value approaches. The experimental data generates a ground truth landscape matrix, while the model produces a corresponding predicted landscape, enabling direct comparison of mutation effects across the entire sequence space (Fig. 1c) for loss computing. This approach facilitates the simultaneous learning of complex mutational interactions and their collective impact on protein expression. We formulated a landscape-oriented Binary Cross-Entropy (BCE) loss function that operates directly on the constructed landscapes (Fig. 1d). The loss computation is specifically constrained to experimentally validated positions, effectively handling the sparse nature of mutation data while maximizing the utilization of available information. Our approach notably improves training efficiency (Fig. S1), enabling the prediction of all single-site mutations with a single forward/backward pass, bypassing the need for per-mutant latent representation calculations. Additionally, our model does not introduce extra parameters or computational overhead during inference, maintaining the zero-shot predictive capability of the original ESM-2. We will show that the model also exhibits cross-protein generalization, demonstrating the ability to predict unseen mutations across diverse proteins. Overall, our method provides an efficient solution with faster training and inference speeds than conventional models, enhancing the model’s capacity to extract meaningful patterns from limited experimental data.

### 2.2 Performance on High-Throughput Datasets

To systematically evaluate our model’s performance across different scales of protein expression data, we analyzed datasets spanning various protein families.(See more details in Methods) The size distribution of these datasets ranges from small-scale collections with fewer than 200 sequences to large-scale datasets containing over 8,000 sequences (Fig. 2a), providing a comprehensive test bed for model evaluation. We observed that the proportion of negative data generally exceeds that of positive data — an anticipated imbalance, as most DMS datasets rely on random mutagenesis, making beneficial mutations relatively rare. We employed a leave-one-out cross-validation strategy on these datasets, and such imbalance poses an additional challenge to the model’s transferability.

**Fig. 2.**
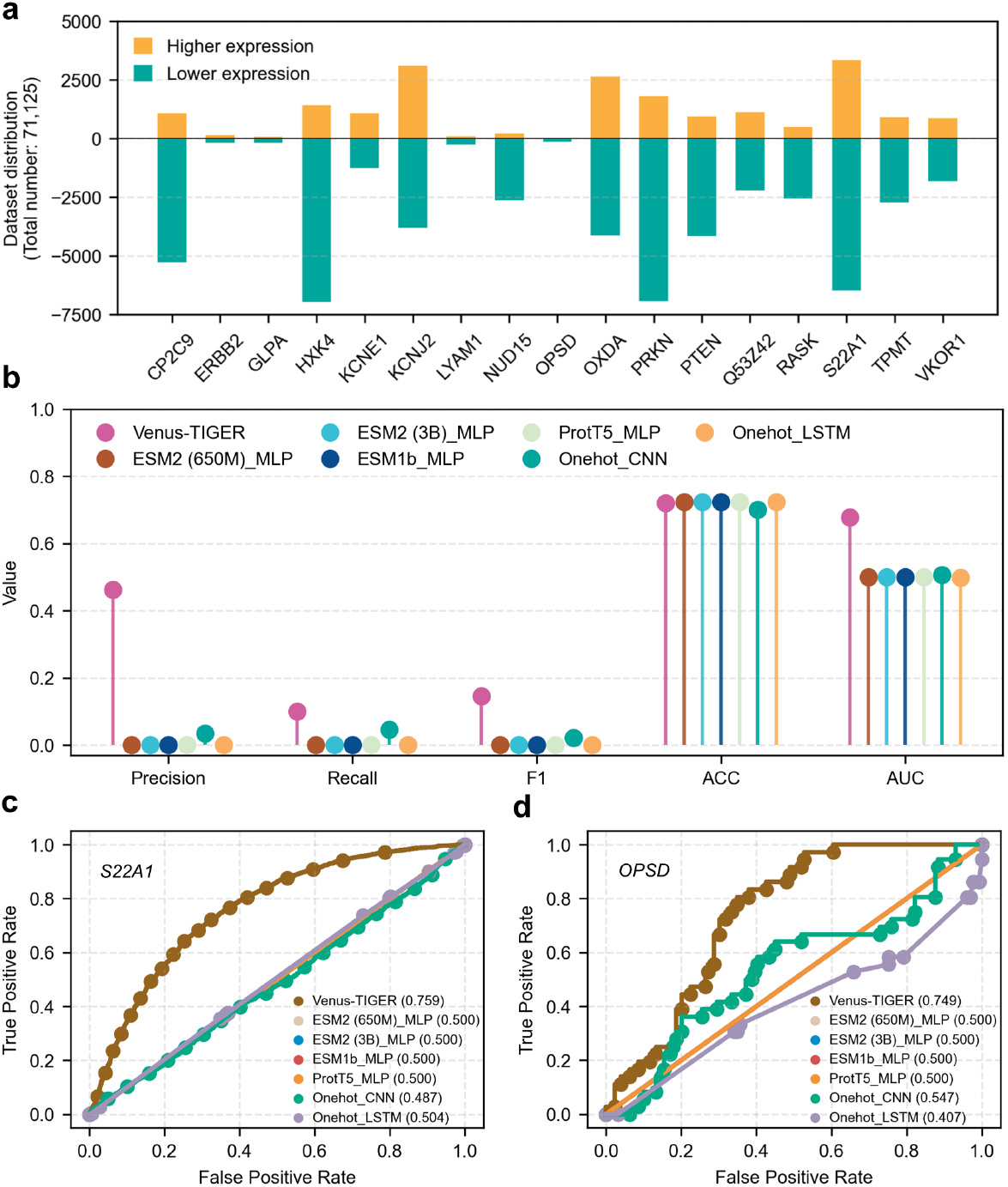
Performance on High-throughput Datasets. **a.** Dataset distribution: Overview of protein expression datasets used in this study, showing the size distribution of different protein families ranging from small datasets (<200 sequences) to large datasets (>8000 sequences). **b**. Comprehensive evaluation metrics (ACC, AUC, F1, Recall, and Precision) of Venus-TIGER and baseline models on high-throughput dataset. ROC curves demonstrating model performance on **c**. the largest dataset (S22A1, n=9,803) and **d**. the smallest dataset (OPSD, n=165) in our collection.

Venus-TIGER demonstrated robust performance across this diverse range of datasets. To comprehensively evaluate our model, we conducted a comparative analysis using multiple evaluation metrics, including Accuracy (ACC), Area Under the Receiver Operating Characteristic Curve (AUC), F1 score, Recall, and Precision. Across all these metrics, Venus-TIGER consistently outperformed baseline approaches, such as other PLMs combined with regression techniques and traditional deep learning methods (Fig.2b).

Notably, in the Precision, Recall, and F1 score metrics, baseline methods yielded scores approaching zero, whereas Venus-TIGER achieved notably higher values. This stark contrast highlights the limitations of conventional approachs in predicting protein expression. The poor performance of these baseline methods can be attributed to their inadequate transferability and scalability in capturing the complex, multifaceted landscape of protein expression. Additionally, sequence-based regression approaches often struggle with imbalanced datasets and fail to effectively identify critical mutations that enhance expression, leading to low Precision and Recall. In contrast, the landscape-based training strategy in Venus-TIGER enables it to simultaneously model the entire expression landscape, thereby overcoming the transferability challenges inherent in baseline methodologies.

Furthermore, we systematically evaluated the performance of different methods on two representative datasets that represent the extremes of our testing spectrum: S22A1 [26] (*>* 8, 000 sequences) and OPSD [27] (*<* 200 sequences) dataset as illustrated in Fig. 2c and d. Receiver Operating Characteristic (ROC) curve analysis revealed that Venus-TIGER consistently outperformed all baseline methods across both scales. In the large-scale S22A1 dataset, Venus-TIGER achieved an AUC of 0.759, while traditional approaches such as Onehot_CNN and Onehot_LSTM performed near random chance (AUC=0.487 and 0.504, respectively). More remarkably, when tested on the small-scale OPSD dataset, Venus-TIGER maintained robust performance with an AUC of 0.749, whereas conventional methods showed notable performance degradation, with Onehot_LSTM achieving only an AUC of 0.407. Various MLP-based approaches, including ESM2, ESM1b, and ProtT5, exhibited consistent but moderate performance (AUC ≈ 0.5) across both datasets. These results demonstrate the remarkable scalability and robustness of Venus-TIGER, particularly its ability to maintain high predictive performance even with limited training data.

### 2.3 Performance on Low-Throughput Dataset of Enzyme (T7 RNAP)

To validate our model’s effectiveness on low-throughput experimental data, we applied Venus-TIGER to predict expression of T7 RNAP mutations. T7 RNAP is one of the workhorses in molecular biology and biotechnology [28–30]. It plays a pivotal role in in vitro transcription and has substantial industrial value, making the study of its expression highly significant [31–33]. T7 RNAP comprises 883 amino acids and is organized into four domains [34]. We conducted a comprehensive structural analysis of mutation effects across different functional domains of T7 RNAP. The spatial distribution of expression-altering mutations was visualized in the N-terminal domain (NTD) (Fig. 3a), thumb domain (Fig. 3b), finger domain (Fig. 3c), and palm domain (Fig. 3d), where red and blue spheres represent mutations that increase and decrease expression, respectively. In our limited dataset, mutations in the NTD, thumb domain, and finger domain were more frequently associated with increased expression compared to decreased expression, whereas mutations in the palm domain often led to relatively lower expression. Despite the palm domain being the key catalytic domain, catalytic activity does not necessarily correlate directly with expression. This finding highlights the complexity of predicting expression from sequence or structural features alone.

**Fig. 3.**
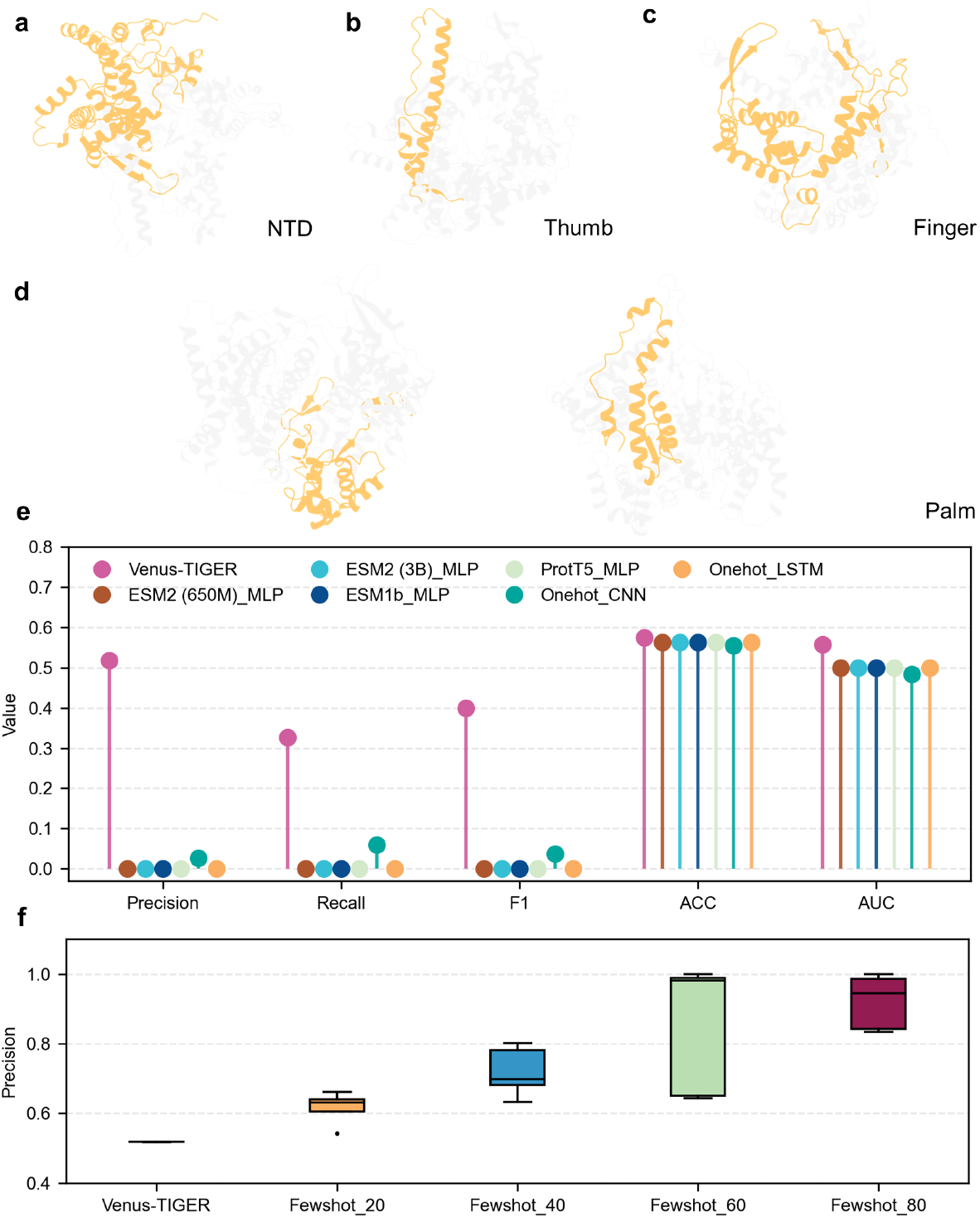
Performance evaluation of Venus-TIGER on T7 RNAP expression prediction. **a-d** Structural visualization of mutation effects on T7 RNA polymerase expression across different domains: **a.** N-terminal domain (NTD), **b**. thumb domain, **c**. finger domain, and **d**. palm domain. Mutations indicated increase expression colored in red, while mutations indicated decrease expression colored in blue. **e**. Comparative performance metrics (Precision, Recall, F1 score, Accuracy, and AUC) of Venus-TIGER and other models on T7 RNAP expression prediction. **f**. Few-shot learning performance showing model predictions with increasing proportions (0%, 20%, 40%, 60%, and 80%) of T7 RNAP experimental data added to the original training dataset.

Venus-TIGER demonstrated superior performance in predicting T7 RNAP expression compared to benchmark models. We evaluated the model using multiple performance metrics, including Precision, Recall, F1 score, ACC, and AUC (Fig. 3e). Our model consistently outperformed existing approaches across all metrics, validating the effectiveness of our landscape-based training strategy on enzyme expression prediction. This is consistent with the results from the previously mentioned DMS dataset, demonstrating the robustness of Venus-TIGER.

Moreover, we conducted an incremental training analysis to investigate the model’s learning efficiency with limited experimental data. We systematically added different proportions (0%, 20%, 40%, 60%, and 80%) of T7 RNAP experimental data to the original training dataset (Fig. 3f). Beyond the performance in zero-shot, the performance of Venus-TIGER shows consistent improvment with increasing amounts of T7 RNAP-specific training data. This progressive enhancement in performance with minimal experimental data underscores Venus-TIGER’s potential for practical applications where extensive mutation datasets may be unavailable or cost-prohibitive to generate, while maintaining its robust baseline performance as demonstrated in the zero-shot scenario.

### 2.4 Performance on Low-Throughput Dataset of Antibody (VHH)

To further evaluate our model’s generalization capability across different protein families, we applied Venus-TIGER to predict expression of mutations in VHH, a single-domain antibody [35, 36]. Due to its compact size, monomeric nature, robust structure, and ease of modification, VHH has become a valuable tool in medical research and clinical antibody drug development [37, 38]. Moreover, it has been utilized as an affinity ligand for the selective purification of various biopharmaceuticals [39–41]. As an antibody, the VHH requires robust expression during production to ensure sufficient yields for both research and potential therapeutic uses. This particular VHH variant consists of 142 amino acids and is composed solely of a single heavy chain variable domain, reflecting its streamlined structure compared to conventional antibodies.

We systematically analyzed mutation effects across the entire VHH sequence, dividing it into three regions for detailed examination: residues 1-50 (Fig. 4a), residues 51-100 (Fig. 4b), and residues 101-142 (Fig. 4c). Based on the current data, variants exhibiting increased expression and those exhibiting decreased expression appear to be relatively evenly distributed, although there is an overall higher proportion of variants with reduced expression. Furthermore, no clear correlations have emerged between specific sequence features or structural locations and expression. These findings suggest that the factors influencing expression are multifaceted, potentially requiring larger datasets and more advanced analytical methods to fully elucidate the underlying mechanisms governing expression changes.

**Fig. 4.**
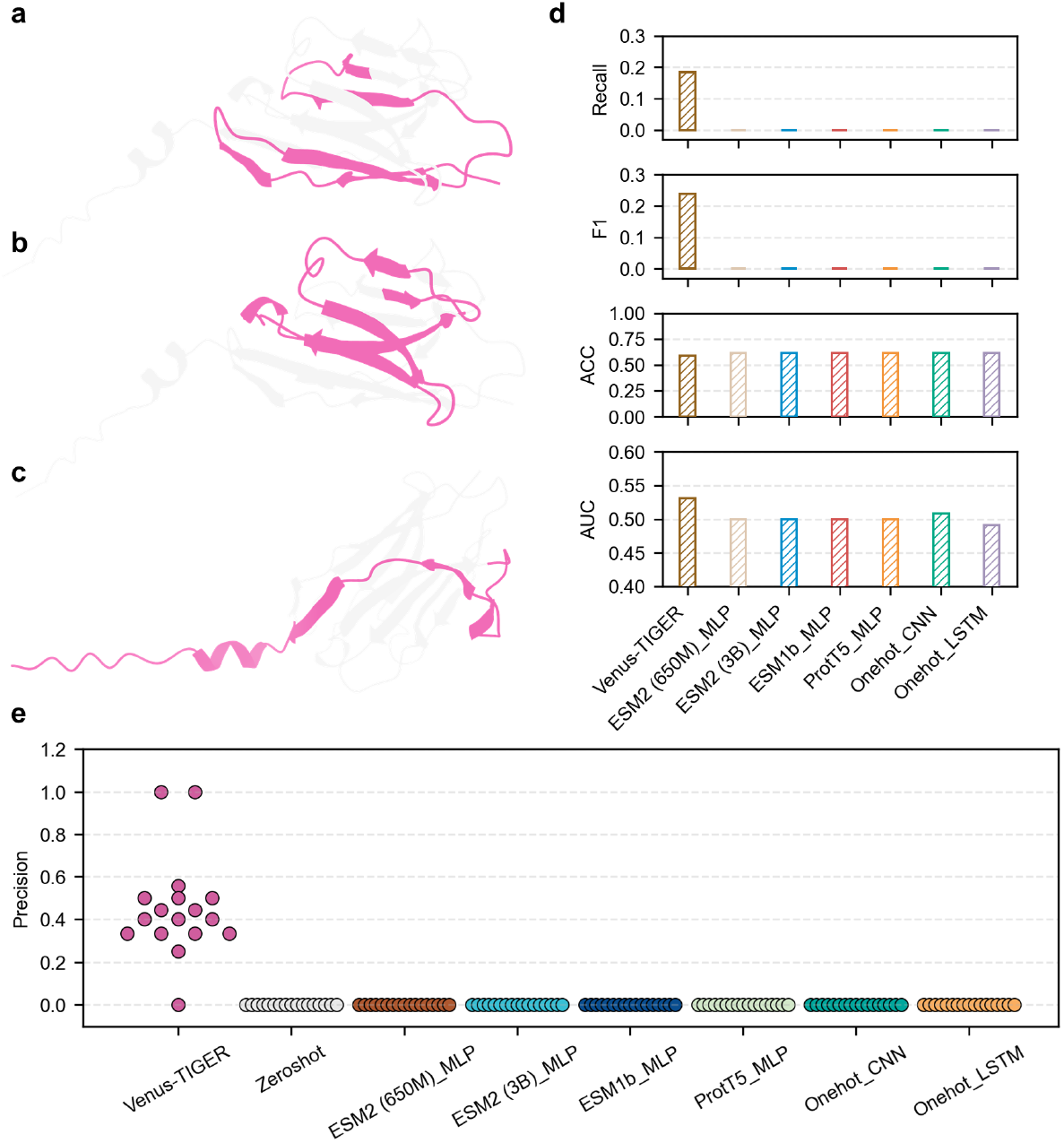
Performance evaluation of Venus-TIGER on VHH expression prediction. Structural visualization of mutation effects on VHH antibody across different sequence regions: **a**. residues 1-50, **b**. residues 51-100, and **c**. residues 101-142. Mutations indicated increase expression colored in blue, while mutations indicated decrease expression colored in green. **d**. Comparative performance metrics (Recall, F1 score, Accuracy, and AUC) between Venus-TIGER and benchmark models. **e**. Comprehensive precision comparison across all 17 validation-test set pairs in leave-one-out cross-validation, showing individual performance of Venus-TIGER and benchmark models.

Venus-TIGER demonstrated exceptional predictive performance for VHH expression compared to benchmark approaches. Quantitative evaluation across multiple metrics revealed substantial improvements over baseline methods (Fig. 4d), with notably higher Recall, F1 score, ACC, and AUC values.

We next implemented a leave-one-out cross-validation framework across 17 different validation-test set combinations to rigorously assess model stability and reliability. As shown in Fig. 4e, Venus-TIGER exhibited consistently superior precision, while all comparative methods - including zero-shot predictions, protein language models, and traditional deep learning approaches - failed to demonstrate meaningful predictive capability, yielding precision scores of zero. This stark contrast in performance underscores the robust generalization capability of our matrix-based training strategy.

The comprehensive evaluation on VHH antibody, together with our previous results on T7 RNAP, demonstrates Venus-TIGER’s broad applicability and reliable performance across diverse protein families, establishing it as a versatile tool for protein expression prediction.

### 2.5 Identifying Expression-Associated Features

To investigate the fundamental determinants of protein expression, we conducted comprehensive statistical analyses of sequence and structural features across on both DMS datasets and T7 RNAP predictions. Assessed preferences in the types of amino acids associated with high or low expression revealed consistent patterns between high-throughput experimental data (gray) and T7 RNAP predictions (pink) (Fig. 5a), particularly for residues with strong positive or negative associations. Specifically, we observed that the amino acids alanine (Ala), threonine (Thr), and methionine (Met) are more frequently associated with high expression levels, whereas proline (Pro), arginine (Arg), tryptophan (Trp), tyrosine (Tyr), glycine (Gly), and lysine (Lys) exhibit a preference for low expression. Furthermore, statistical analysis indicates that mutations to basic amino acids (histidine [His], arginine [Arg], and lysine [Lys]) are unfavorable for high expression, while mutations to acidic amino acids (aspartic acid [Asp] and glutamic acid [Glu]) tend to positively influence expression. Additionally, aromatic amino acids (Tyr, Trp, and phenylalanine [Phe]) are more often linked to low expression. These phenomena show similar tendencies across both datasets, validating our model’s capacity to capture intrinsic sequence-expression relationships.

**Fig. 5.**
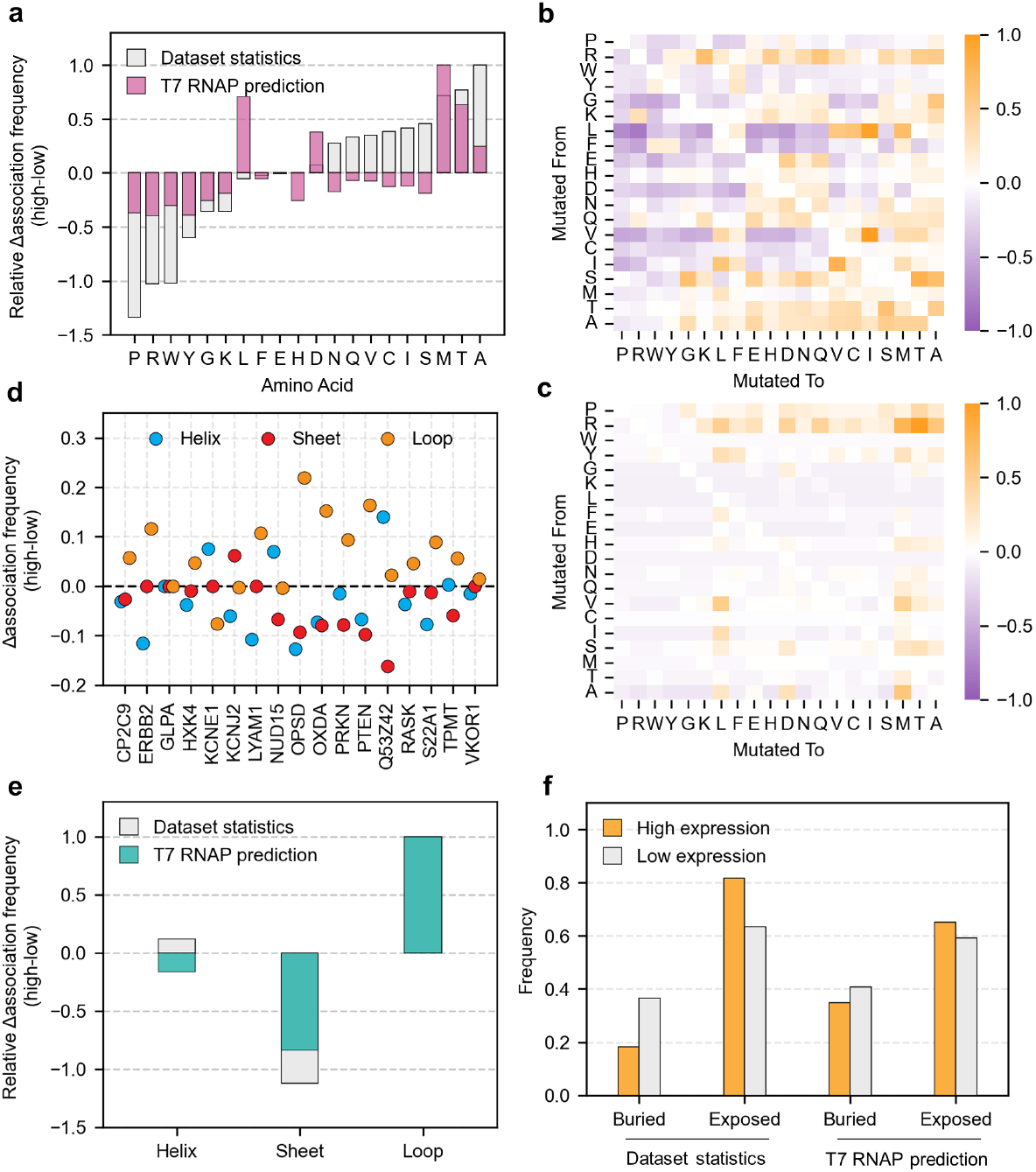
Statistical analysis of amino acid preferences and structural features associated with protein expression. **a.** Expression-associated amino acid preferences comparing high-throughput datasets (gray) and T7 RNAP predictions (pink). **b**,**c**. Analysis of amino acid pair correlations with expression in **b**. high-throughput datasets and **c**. T7 RNAP single-site mutations. **d**. Secondary structure preferences across 17 high-throughput datasets, showing relationships between mutation positions (helix, sheet, loop) and expression. **e**. Comparative analysis of secondary structure preferences between high-throughput datasets (gray) and T7 RNAP predictions (green). **f**. Correlation between residue exposure and expression, comparing high-throughput datasets (left panel) and T7 RNAP predictions (right panel).

The amino acid pair correlation analysis unveiled complex interaction patterns influencing expression levels. In the heatmap of mutational pairs (Fig. 5b), both the horizontal and vertical axes are arranged according to the statistical frequency of each amino acid’s association with low expression (on the left) to high expression (on the right). The color scale indicates purple for pairs correlated with low expression and yellow for pairs correlated with high expression. A higher density of purple is evident in the left region, while yellow clusters dominate the right region, implying that substituting amino acids associated with low expression for those linked to high expression often leads to elevated expression. The heatmap derived from high-throughput datasets (Fig. 5b) revealed distinct mutational preferences, with certain amino acid substitutions consistently associated with expression changes. Notably, these patterns were independently reproduced in our T7 RNAP single-site mutation predictions (Fig. 5c), suggesting mostly similar molecular mechanisms governing protein expression at individual sites across diverse protein families.

Secondary structure analysis revealed distinct patterns of mutation effects across different structural elements. Analysis across 17 high-throughput datasets showed varying impacts of mutations in helix, sheet, and loop regions (Fig. 5d), with loop regions showing predominantly positive association frequencies, whereas sheet regions were frequently associated with low expression. The aggregated analysis (Fig. 5e) demonstrated remarkable concordance between high-throughput experimental data (gray) and T7 RNAP predictions (green), with mutations in loop regions consistently associated with enhanced expression, while mutations in sheet structures typically showed negative effects.

By examining the relationship between residue exposure and expression, we found that mutations at exposed positions more frequently led to higher expression, whereas buried positions were predominantly correlated with lower expression. These results were consistently observed in both experimental data and T7 RNAP predictions (Fig. 5f), indicating that exposed residues are generally more tolerant to mutations than their buried counterparts. In particular, exposed residues exhibited a higher frequency of expression-altering mutations compared to buried residues, suggesting that residue accessibility is a fundamental determinant of how mutations affect protein expression. These comprehensive analyses not only validate Venus-TIGER’s predictive capabilities but also provide insights into the structural and sequence determinants of protein expression.

### 2.6 Correlations among Activity, Stability and Expression

To understand the complex interplay between different protein properties, we conducted a systematic comparison of sequence and structural preferences across expression (n=71,125), stability (n=143,965), and activity (n=65,887) using DMS datasets. Our analysis revealed both shared and distinct patterns among these properties.

From the perspective of amino acid preferences, we observed that activity and expression showed similar patterns in their amino acid preference, with residues such as Ala, Thr and Met showing consistently positive associations, while Pro, Arg, and Trp exhibited negative correlations for both properties (Fig.6a,g). However, stability patterns (Fig.6d) displayed distinct characteristics, particularly evident in residues like Arg, Thr and Ser, which showed opposing effects compared to expression and activity. Notably, several residues (Ala, Pro, Met, Cys, and Ile) showed consistent effects across all three properties (Fig. 6a,d,g), suggesting their fundamental roles in protein function. Analysis of secondary structure preferences revealed complex relationships among these properties. Expression and stability demonstrated similar trends in their association with secondary structural elements, while activity showed distinct preferences. A consistent pattern emerged where mutations in loop regions were associated with enhanced performance across all properties, while mutations in sheet regions generally showed negative impacts (Fig. 6b,e,h). The impact of residue exposure emerged as a unifying principle across all three properties. Exposed residues consistently showed higher frequencies of beneficial mutations compared to buried positions (Fig. 6c,f,i). This uniform pattern suggests that residue accessibility plays a fundamental role in determining protein properties, possibly by influencing protein folding and molecular interactions.

**Fig. 6.**
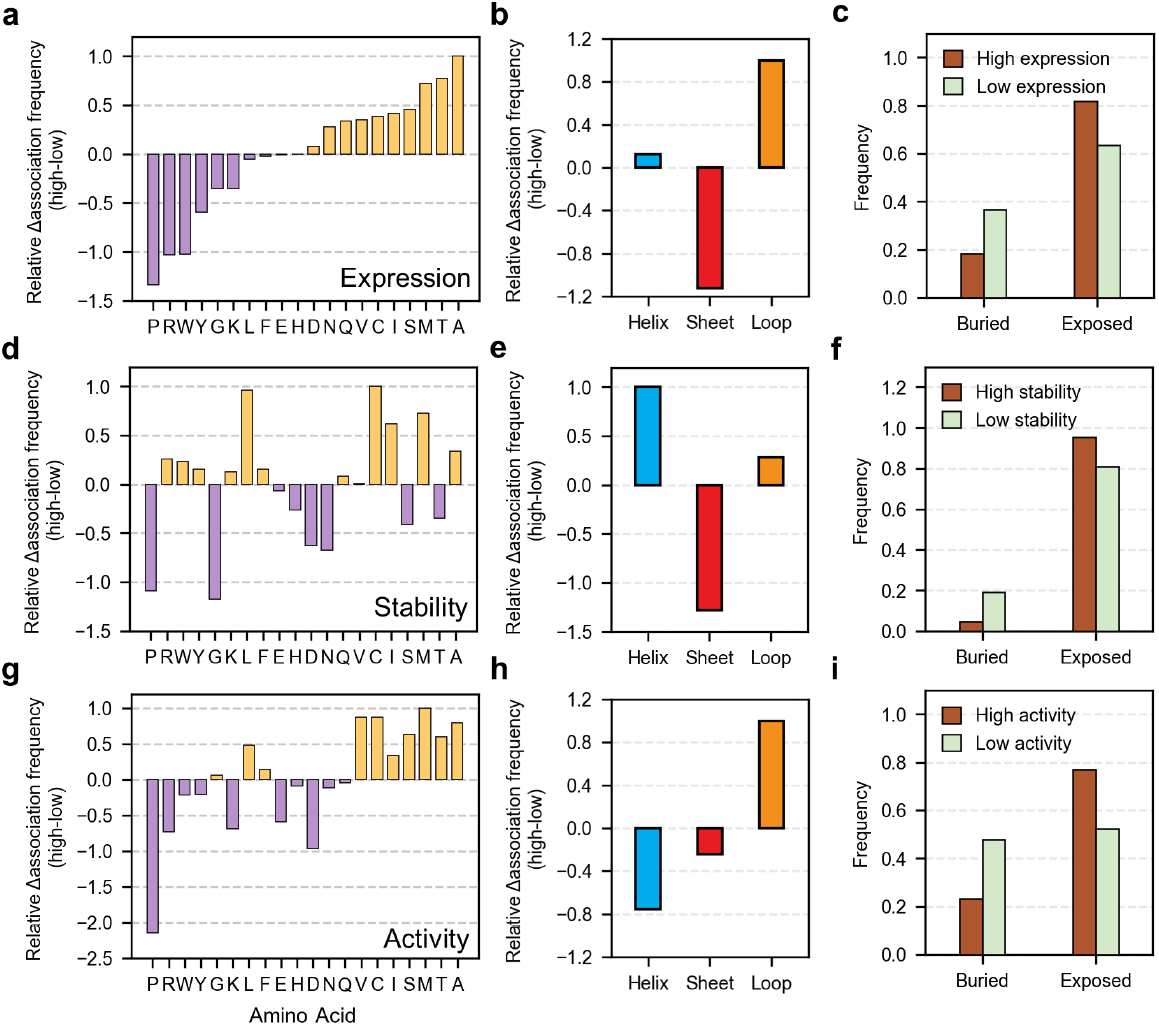
Comparative analysis of sequence and structural preferences across protein expression, stability, and activity. **a-c.** Analysis of expression-associated features: **a**. amino acid preferences, **b**. secondary structure preferences (helix, sheet, loop), and **c**. relationship between residue exposure and expression. **d-f**. Analysis of stability-associated features: **d**. amino acid preferences, **e**. secondary structure preferences, and **f**. relationship between residue exposure and stability. **g-i**. Analysis of activity-associated features: **g**. amino acid preferences, **h**. secondary structure preferences, and **i**. relationship between residue exposure and activity.

These comprehensive analyses reveal the intricate interplay between different protein properties and their structural determinants. The identified patterns of commonality (particularly in residue exposure effects) and distinction (notably in secondary structure preferences) provide mechanistic insights into the underlying mechanisms governing protein function.

## 3 Discussion

In this study, we have developed a deep learning framework for predicting and understanding expression-governing residues. We incorporated a PLM with a top-layer regression model instead of adding external MLP layers, improving compatibility with the feature extraction backbone. The landscape loss function processes entire datasets simultaneously, reducing training iterations and improving model transferability. This approach demonstrates consistent performance across high-throughput datasets and transfers effectively to low-throughput experimental data, as validated on two distinct protein families: an enzyme (T7 RNAP) and an antibody (VHH). The model’s performance improves progressively with increasing proportions of T7 RNAP data incorporated into the original training dataset, demonstrating efficient adaptation to new protein families. This capability addresses a critical challenge in protein engineering, where experimental data acquisition is resource-intensive, while maintaining the benefits of pre-trained features.

Our comprehensive statistical analyses have revealed patterns in amino acid preferences, amino acid pair interactions, secondary structure contexts, and exposure preferences across high-throughput datasets. Remarkably, these patterns were largely reproduced in our analysis of T7 RNAP saturation mutagenesis predictions, with minor variations attributable to protein-specific characteristics. This consistency validates our model’s ability to capture fundamental biophysical features that influence protein expression, demonstrating both its learning capacity and transferability while providing mechanistic insights from a biophysical perspective.

Furthermore, our comparative analysis of stability, activity, and expression preferences has uncovered intriguing relationships among these protein properties. Activity and expression exhibited similar amino acid preferences, whereas stability displayed some distinct patterns. Nevertheless, all three properties demonstrated consistent preferences for specific amino acids (Ala, Pro, Met, Cys, and Ile) in their contributions to expression. While expression and stability shared similar secondary structure preferences, activity manifested distinct patterns. Notably, all three properties consistently associated loop regions with enhanced performance and sheet regions with reduced performance. Furthermore, exposed residues uniformly correlated with improved performance across all three properties, suggesting a fundamental role of solvent accessibility in protein function. These shared and distinct patterns offer valuable insights into the interconnected nature of these properties and their underlying biophysical mechanisms.

This work advances both computational methodology and biological understanding of protein expression. The landscape-based training strategy and modified model architecture enable efficient prediction across diverse protein families. Statistical analyses of high-throughput datasets and model predictions reveal consistent patterns in sequence-structure-function relationships, providing insights into the mechanisms underlying protein expression, stability, and activity. These findings contribute to the fundamental understanding of protein properties and their applications in protein engineering.

## 4 Methods

### 4.1 Model Details

#### 4.1.1 Model Architecture

The architecture of Venus-TIGER is the same as that of ESM-2 (650M Version) [15], which is a transformer equipped with rotary position embedding [42] and pre-layer normalizations. Venus-TIGER accepts protein sequences *S* = {*s*_1_, *s*_2_, …, *s*_*L*_}, where *s*_*i*_ represents the one-hot encoding of amino acid at position *i* in the sequence, and *L* is the sequence length. After passing through an embedding layer and 33 Transformer encoder layers, the model outputs contextualized embeddings ***H*** = {***h***_1_, ***h***_2_, …, ***h***_*L*_}, where ***h***_*i*_ ∈ ℝ^*d*^ (*d* = 1280) represents the latent presentation of amino acid *s*_*i*_ considering its sequence context. Finally, the model uses a multi-layer perceptron (MLP) to map the embeddings *h*_*i*_ back to amino acid space 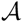. For each position *i*, the MLP returns a probability distribution over all possible amino acids:

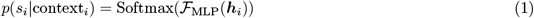

where 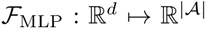 is the MLP layer; context_*i*_ = {***h***_1_, …, ***h***_*i*−1_, ***h***_*i*+1_, …, ***h***_*L*_} is the sequence context excluding position *i* and *p*(*s*_*i*_|context_*i*_) is the probability distribution over amino acid at position *i*.

#### 4.1.2 Mutation Effect Prediction

One of the most common applications of the above model is to predict the effect of protein mutations [15, 19, 43]. Formally, a single-site mutation can be represented as a tuple 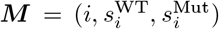, where *i* is the position of the mutation in the sequence, 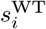is the wild-type amino acid at position *i* and 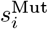 is the mutant amino acid replacing 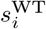. For example, a mutation replacing the amino acid at position 10 from Alanine (A) to Glycine (G) is represented as *M* = (10, *A, G*). For a wild-type amino acid 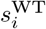 at position *i*, the log-likelihood is:

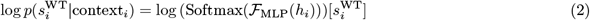

Similarly, for a mutant amino acid 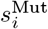, the log-likelihooid is:

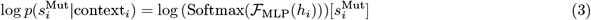

Then, the mutation effect can be estimated by the log-likelihood difference [15, 16]:

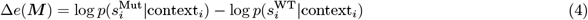

#### 4.1.3 Binary Mutation Effect Prediction

We can also classify the effect of a mutation as binary, as it determines whether a mutation is positive or negative. This binary classification helps identify critical positions and amino acids essential for protein expression level, stability, or other features. In general, we define a binary zero-shot mutation effect prediction method based on the log-likelihood difference:

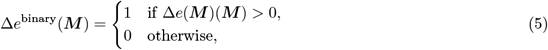

where Δ*e*^binary^(*M*) = 1 indicates a positive mutation and Δ*e*^binary^(*M*) = 0 indicates a negative mutation.

#### 4.1.4 Parallel Landscape Computation Using Matrix Operations

To efficiently perform back-propagation and forward computation, we utilize matrix operations to facilitate parallel computations. This approach allows us to generate a saturated mutation fitness landscape during one forward computation pass. Let 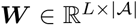 denote the one-hot encoding matrix of the wild-type sequence. We can compute the effect for all possible mutations in matrix operation:

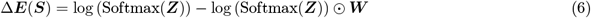

where *Z* = ℱ_MLP_(*H*) is the logits, ⊙ represents element-wise multiplication, and Softmax(*Z*) is the probabilities for all amino acids at each position. For binary mutation effect prediction, we can apply the Equation 5 to obtain the binary mutation effect matrix Δ***E***^*binary*^(***S***).

#### 4.1.5 Training Set Construction and Loss Function

In our settings, the training set includes multiple proteins, each with its own set of measured mutations. In practice, these datasets are often not saturated, meaning that only a subset of all possible single-site mutations has been experimentally measured for each protein. To train on such non-saturated mutation data, we construct a masked ground truth matrix for each protein (Fig. 1c). A PyTorch-style pseudo code snippet for constructing training set is provided in Algorithm 1 of the Supplementary Information. Given a dataset of *N* proteins, where each protein *κ* has a sequence 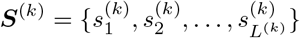 with length *L*^(*κ*)^, and a set of measured mutations 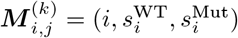, we proceed as follows:

1. **Masking Matrix Construction**. For each protein *κ*, we define a masking matrix 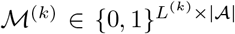. Each item 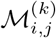 indicates whether the mutation 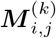 is present in the measured data:

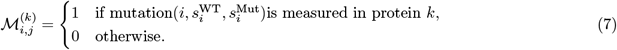
2. **Ground Truth Matrix Definition**. Correspondingly, for each protein *k*, we define a binary ground truth matrix 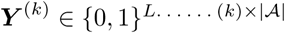, where:

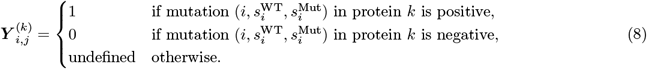 Only the items where 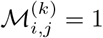 are defined and utilized in the loss computation.
3. **Binary Cross-Entropy Loss Function**. To train the model, we employ a binary cross-entropy (BCE, Fig. 1d) loss function that considers only the observed mutations across all proteins, as indicated by the masking matrices ℳ ^(*κ*)^. The total loss ℒ is defined as:

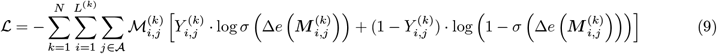

where *σ* denotes the sigmoid function, and

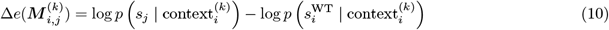

is the log-likelihood difference for mutation 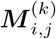 in protein *κ*, and 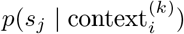 is the predicted probability of amino acid *s*_*j*_ at position *i* in protein *κ*, as defined in Equation 1.

Furthermore, by utilizing the parallel matrix operations described in Equation 6, training a batch of protein mutation datasets requires only one forward and backward pass, notably improving computational efficiency. A PyTorch-style pseudo code snippet for computing loss is listed in Supplementary Information Algorithm 2.

### 4.2 Collection of the Training Data

The training datasets are initially taken from ProteinGym [43], including 17 different proteins and a total of 71,125 sequences. The 17 proteins are CP2C9, ERBB2, GLPA, HXK4, KCNE1, KCNJ2, LYAM1, NUD15, OPSD, OXDA, PRKN, PTEN, Q53Z42, RASK, S22A1, TPMT, and VKOR1, expressed in bacteria, fungi, and mammalian cells. For each protein, we selected the expression scores of the wild type as thresholds and performed binarization to classify the variants into two categories.

### 4.3 In-Silico Experiments

#### 4.3.1 Details of Venus-TIGER

##### High-Throughput DMS Datasets Evaluation

To comprehensively evaluate our model’s performance on high-throughput DMS datasets, we implemented a systematic cross-validation strategy across 17 DMS datasets. For each experiment, we selected one dataset as the validation set and another as the test set, while the remaining 15 datasets served as the training set. This resulted in 272 unique validation-test combinations. The training process utilized a batch size of 1 and was conducted for a maximum of 500 epochs. We employed the Adam [44] optimizer with a learning rate of 1*e*^−4^ and Binary Cross-Entropy (BCE) as the loss function. To prevent overfitting, we implemented early stopping with a patience of 20 epochs, monitoring the validation loss. Model performance was evaluated using multiple metrics, including Precision, Recall, F1 Score, ACC, and AUC, with a classification threshold of 0.5. The final results were obtained by averaging these metrics across all 272 validation-test combinations.

##### Low-Throughput DMS Datasets Evaluation

The low-throughput datasets were generated by experimentally assessing the expression of mutants predicted in our previous work [19, 45, 46]. In the current investigation, these datasets are employed exclusively to test and validate the effectiveness of our model in predicting protein expression. For the low-throughput DMS dataset evaluation (T7 RNAP and VHH), we modified our experimental design while maintaining consistent training parameters. In this setup, we used 16 high-throughput DMS datasets as the training set and selected one remaining high-throughput dataset as the validation set, while consistently using the low-throughput DMS dataset as the test set. This process was repeated 17 times, rotating through each high-throughput dataset as the validation set. The training protocol remained identical to the high-throughput evaluation, with a batch size of 1, maximum 500 epochs, learning rate of 1*e*^−4^, and BCE loss function. Early stopping was similarly implemented with 20 epochs patience, monitoring validation loss. The same evaluation metrics (Precision, Recall, F1 Score, ACC, and AUC) were calculated using a classification threshold of 0.5, and final results were averaged across all 17 validation set combinations to assess the model’s generalization ability on low-throughput data.

##### Protein Sequences Diversity and Similarity Analysis

To ensure that our model can generalize dissimilar proteins to those in the training set, we performed a series of visual and quantitative similarity analyses. The results show that the wild-type sequences collected from the 19 datasets, including the ProteinGym dataset and two validation datasets, T7 RNAP and VHH, exhibit a wide distribution. This is reflected not only in the distances based on embeddings but also in the distances based on sequence alignment, as shown in Fig. S2, Fig. S3, and Fig. S4 in Supplementary Information. First, Fig. S2 presents a two-dimensional UMAP [47] projection of the ESM2-650M derived sequence embeddings. The broad, scattered arrangement of points indicates that the proteins are well-distributed in the latent space without forming tight, overlapping clusters. Next, Fig. S3 shows the distribution of pairwise sequence similarities, calculated via cosine similarity [48] (panel a) and L2 distance [49] (panel b). Here, most sequence pairs exhibit low similarity, suggesting a diverse set of protein variants. Finally, Fig. S4 employs the Needleman-Wunsch alignment algorithm [50] to generate a pairwise similarity matrix (panel a) and a histogram of alignment scores (panel b). Again, the results show that most sequences in the dataset share limited homology (< 30%). Taken together, these analyses demonstrate that our dataset includes a wide range of protein sequence variation, which not only ensures uniform coverage of protein space but also demonstrates the capacity of our model to generalize to proteins with minimal similarity to those seen in training.

#### 4.3.2 Baseline Model

##### Language Models

To benchmark our model’s performance, we first implemented several state-of-the-art protein language models as feature extractors. These include ESM-2 (both 650M and 3B parameter variants), ESM-1b, and ProtT5, representing different scales of pre-trained models with parameters ranging from 650M to 3B [15, 17, 18]. For each language model, we extracted sequence embeddings and processed them through a lightweight MLP classifier implemented in PyTorch Lightning. The classifier consists of two linear layers: the first layer transforms the input embeddings to a hidden dimension of 128 with ReLU [51] activation, followed by a second linear layer that produces the final single-neuron output. This output is then passed through a sigmoid activation function for binary classification. The MLP classifier was trained using Binary Cross-Entropy loss with an Adam optimizer (learning rate = 1*e*^−4^), leveraging the rich protein representations learned by these pre-trained language models while maintaining computational efficiency.

##### Traditional Deep Learning Models

We also implemented two traditional deep learning architectures that directly process one-hot encoded amino acid sequences. The CNN model employs a hierarchical structure beginning with a convolutional layer followed by ReLU activation and max-pooling operations [52]. This is followed by a second convolutional layer with ReLU activation, an adaptive max-pooling layer, and a flattening operation. The flattened features are then processed through a linear layer with ReLU activation, culminating in a final linear layer for classification output. In contrast, the LSTM model adopts a more straightforward architecture, where one-hot encoded sequences are processed through LSTM layers, followed directly by a linear layer for classification [53]. Both architectures were trained end-to-end and evaluated using the same protocol as our proposed model.

For high-throughput datasets, we implemented the 272 validation-test combinations cross-validation strategy, while for low-throughput dataset evaluation, we rotated through 17 validation sets with a fixed test set (T7 RNAP or VHH). Performance metrics including Precision, Recall, F1 Score, ACC and AUC were calculated and averaged across all validation-test combinations, ensuring fair comparison across different architectural approaches.

### 4.4 Wet-Lab Experiments

#### 4.4.1 Protein Expression and Purification of T7 RNAP

The T7 RNAP gene was cloned into the pQE-80L expression vector and transformed into *E. coli* BL21(DE3) cells. The cells were cultured in LB media until reaching an OD600 of approximately 0.6-0.8, followed by induction with 1 mM IPTG for a 6-hour growth period at 37 °C. After collection, the bacteria were resuspended in a binding buffer (50 mM Tris-HCl pH 8.0, 300 mM NaCl, 3 mM imidazole, 0.1 mM EDTA) and lysed via sonication. The resulting lysate underwent centrifugation at 4 °C and 12,000 rpm for 30 minutes. The lysate was then applied to a Ni-NTA gravity column and washed with a washing buffer (50 mM Tris-HCl pH 8.0, 300 mM NaCl, 10 mM imidazole, 0.1 mM EDTA, 10% glycerol). Elution was performed using an elution buffer (50 mM Tris-HCl pH 8.0, 300 mM NaCl, 250 mM imidazole, 0.1 mM EDTA, 10% glycerol). Concentration was achieved using a final ultrafiltration buffer (50 mM Tris-HCl pH 8.0, 100 mM NaCl, 0.1 mM EDTA), and the T7 RNAP was diluted with a storage buffer (50 mM Tris-HCl pH 8.0, 100 mM NaCl, 0.1 mM EDTA, 1 mM DTT, 75% glycerol). [19, 54] The protein yield was quantified by NanoDrop (Thermofisher). We selected the expression scores of the wild type as thresholds and performed binarization to classify the variants into two categories.

#### 4.4.2 Protein Expression and Purification of VHH

The gene of the VHH was cloned into the pET29a plasmid. The expression plasmid was transformed into *E. coli* BL21(DE3) cells. A single colony of each recombinant E. coli strain was inoculated into 30 mL LB medium with 50 ug/mL kanamycin for seed culture at 37 °C for 12-16 h. The seed culture was transferred 10 ml to 1L LB medium with 50 ug/ml kanamycin at 37 °C 220rpm until the OD600 value get 0.6-0.8. The culture was cooled to 16 °C and then induced with 0.5 mM IPTG for 20-24 h at 16 °C. Cells were harvested from the fermentation culture by centrifugation for 30 min at 4,000 rpm, and the cell pellets were collected for later purification. The cell pellets were resuspended in buffer A (20 mM Na2HPO4 and NaH2PO4, 0.5 M NaCl, pH 8.0) and then and lysed via ultra sonification. The lysates were centrifuged for 30 min at 12,000 rpm at 4 °C, after which the supernatants were subjected to Ni-NTA affinity purification with elution buffer (20 mM *Na*_2_*HPO*_4_ and *NaH*_2_*PO*_4_, 0.5 M NaCl, 250 mM imidazole, pH 8.0). The purity of the fractions obtained was analyzed using SDS-PAGE. The fractions containing the purified target protein were combined and desalted using an ultrafiltration unit. The purified protein was then concentrated and stored in buffer A with 10% glycerol at a temperature of −80°C. [19, 45, 46] The protein yield was quantified by NanoDrop (Thermofisher). We selected the expression scores of the wild type as thresholds and performed binarization to classify the variants into two categories.

### 4.5 Statistical Analysis

#### 4.5.1 Dataset Processing

We conducted comprehensive statistical analyses on multiple types of protein mutation datasets, including protein expression, stability, and activity data. For each protein in each dataset, we implemented a standardized binary classification approach based on the wild-type protein’s performance. Mutations were classified as beneficial or deleterious by comparing their scores with the wild-type reference score, establishing a consistent framework for cross-dataset analysis. This binary classification strategy enabled unified analysis across different protein properties while maintaining the biological significance of the mutation effects.

The training data of expression include 17 different proteins and a total of 71,125 sequences. The 17 proteins are CP2C9, ERBB2, GLPA, HXK4, KCNE1, KCNJ2, LYAM1, NUD15, OPSD, OXDA, PRKN, PTEN, Q53Z42, RASK, S22A1, TPMT, and VKOR1, expressed in bacteria, fungi, and mammalian cells. [43] The training data of stability include 64 different proteins and a total of 65,887 sequences. The 64 proteins are AMFR, ARGR, BBC1, BCHB, CATR, CBPA2, CBX4, CSN4, CUE1, DN7A, DNJA1, DOCK1, EPHB2, FECA, FKBP3, HCP, HECD1, ILF3, ISDH, MAFG, MBD11, MYO3, NKX31, NUSA, NUSG, OBSCN, ODP2, OTU7A, PIN1, PITX2, PKN1, POLG, PR40A, PSAE, RAD, RBP1, RCD1, RCRO, RD23A, RFAH, RL20, RPC1, RS15, SAV1, SBI, SCIN, SDA, SOX30, SPA, SPG2, SPTN1, SQSTM, SR43C, SRBS1, TCRG1, THO1, TNKS2, UBE4B, UBR5, VG08, VILI, VRPI, YAIA and YNZC expressed in bacteria, fungi, and mammalian cells. [55] The training data of activity include 38 different proteins and a total of 143,965 sequences. The 38 proteins are A0A1I9GEU1, A0A247D711, ADRB2, AICDA, AMIE, ANCSZ, CAS9, CASP3, CASP7, CCDB, D7PM05, ENVZ, GFP, HEM3, KCNE1, KCNH2, KCNJ2, LGK, MET, OTC, OXDA, PAI1, PHOT, PPARG, PTEN, Q6WV13, Q8WTC7, Q837P4, Q837P5, Q59976, RASH, RL40A, RNC, S22A1, SC6A4, SRC, UBE4B, VKOR1 expressed in bacteria, fungi, and mammalian cells. [43]

#### 4.5.2 T7 RNAP Saturation Single-mutant Analysis

For the T7 RNAP analysis, we generated a comprehensive saturation mutagenesis dataset using our model predictions. Each possible single-site mutation was evaluated, and the predicted scores were binarized using a threshold of 0.5 to classify mutations as either beneficial or deleterious. This systematic approach provided a complete landscape of potential mutation effects, enabling detailed analysis of position-specific and amino acid-specific patterns in T7 RNAP.

#### 4.5.4 Amino Acid Analysis

We analyzed both single amino acid and amino acid pair preferences associated with protein properties. For single amino acid analysis, we calculated the frequency of each amino acid type in mutations leading to increased performance (*f*_*high*_) and decreased performance (*f*_*low*_) in expression, stability, and activity separately. For amino acid pair analysis, we computed the frequencies of wild-type to mutant amino acid substitutions associated with expression changes. In both analyses, the delta frequency was calculated as *δf* = *f*_*high*_ − *f*_*low*_ to quantify the association with protein properties. These calculations were performed for all dataset to identify consistent patterns in amino acid preferences.

#### 4.5.4 Structural Analysis

Using PyMOL and Python scripts, we analyzed the relationship between protein structural features and performance (expression, stability, and activity). Secondary structure elements (*α*-helices, *β*-sheets, and loops) were assigned for each residue position, and the delta frequency (*δf* = *f*_*high* −_ *f*_*low*_) was calculated to quantify the association between structural elements and performance. Solvent Accessible Surface Area (SASA) was calculated for each residue [56], with positions classified as exposed (*SASA* > 20Å^2^) or buried. For exposure analysis, we compared the frequencies of high-performance and low-performance mutations in exposed versus buried positions. These analyses were performed separately for expression, stability, and activity datasets.

#### 4.5.5 Training and Inference Speed Analysis

We evaluated the time cost of our model for inference and training. The used machine is equipped with an AMD EPYC 7H12 64-Core Processor and an NVIDIA RTX 4090, running Ubuntu 22.04 LTS and PyTorch version 2.4. We tested both our model and the baseline model on all 71,225 mutant sequences for inference and training times (one epoch). The training time included two parts: the time to generate embeddings and the time to train one epoch. Each test was repeated five times to calculate the average. As shown in Supplementary Information Fig. S1, our model achieved the best efficiency with times of 0.0537s for generating embeddings, 0.40s for training one epoch, and 0.48s for inference. Additionally, we have also listed the number of parameters of all models.

## 5 Data Availability

The data supporting this article are included as part of the Supplementary Information.

## 6 Acknowledgements

We thank the Computational Biology Key Program of Shanghai Science and Technology Commission (23JS1400600), Shanghai Jiao Tong University Scientific and Technological Innovation Funds (21X010200843), and Science and Technology Innovation Key R & D Program of Chongqing (CSTB2022TIAD-STX0017) for their support. We also acknowledge the contributions from the Student Innovation Center at Shanghai Jiao Tong University and Shanghai Artificial Intelligence Laboratory.

## 7 Author Contributions

Fan Jiang, Bozitao Zhong, and Mingchen Li contributed to the conceptualization of the study. Mingchen Li proposed and implemented the model. Fan Jiang and Yuanxi Yu curated the dataset. Fan Jiang and Banghao Wu did the wet-lab experiments. Fan Jiang ran the all in-silico experiments and did the analysis. Liang Hong supervised the project and provided funding. Fan Jiang drafted the manuscript and visualized the data. All authors participated in discussion and revising the manuscript.

## Footnotes

1 https://huggingface.co/facebook/esm2_t33_650M_UR50D

